# Effect of ORL-1 on Cav1.2 calcium channels

**DOI:** 10.64898/2026.07.03.736403

**Authors:** Aiden J. Shaver, Ivana A. Souza, Laurent Ferron, Maria A. Gandini, Gerald W. Zamponi

## Abstract

Cav1.2 is an L-type voltage-gated Ca^2+^ channel (VGCC) that supports Ca^2+^ influx in response to membrane depolarization. Ca^2+^ entering via Cav1.2 alters gene expression, activates Ca^2+^-dependent enzymes and has been implicated in synaptic plasticity. ORL-1 is a Gi/o-coupled G protein-coupled receptor (GPCR) that is expressed in the peripheral and central nervous systems. Both Cav1.2 and ORL-1 are expressed in the hippocampus, where they have been implicated in learning and memory. It is well-documented that ORL-1 interacts with another VGCC, Cav2.2. However, less is known about potential interactions between Cav1.2 and ORL-1. Here, we examine the interplay between Cav1.2 (Cavα1c, Cavα2δ-1, Cavβ1b) and ORL-1 co-expressed in tsA-201 cells by using biochemical, electrophysiological and confocal imaging analysis. Co-immunoprecipitations revealed that ORL-1 independently interacts with Cavα1c and Cavα2δ-1 subunits of the Cav1.2 channel complex. Electrophysiological recordings revealed that co-expression with ORL-1 reduced Cav1.2 peak current density without altering its biophysical properties. Acute perfusion with the ORL-1 receptor agonist nociceptin (1 µM) did not alter Cav1.2 current density. Confocal imaging experiments revealed that ORL-1 significantly decreases Cav1.2 plasma membrane expression by disrupting forward trafficking. Interestingly, ORL-1 did not affect Cav1.2 endocytosis. Overall, our results demonstrate a previously unrecognized interaction between ORL-1 and Cav1.2 that alters Cav1.2 membrane expression without affecting biophysical properties.

## Introduction

Voltage-gated calcium channels (VGCCs) are a family of voltage-sensitive proteins that open in response to membrane depolarization, facilitating calcium influx [4]. In neurons, VGCCs have been shown to play pivotal roles in many functions, including neurotransmitter release and excitation-transcription coupling [13, 62]. They can be classified into high-voltage-activated channels, which are composed of the pore-forming subunit Cavα1 and the auxiliary subunits Cavα2δ and Cavβ, and low-voltage-activated channels, which are comprised of only the Cavα1 subunit [12, 22, 42]. Cav1.2 is a high-voltage-activated calcium channel and a member of the L-type calcium channel family. It is highly expressed throughout the nervous system, including in hippocampal neurons, where it is primarily clustered in dendritic shafts and spines [21, 29, 57]. In the brain, the physiological functions of Cav1.2 range from excitation-transcription coupling to activation of calcium-dependent enzymes and regulation of neuronal plasticity. Cav1.2 has been implicated in various aspects of learning and memory, including memory acquisition [56], maintenance [5, 28, 54], destabilization [40, 58], and extinction [9, 20, 28, 60].

The opioid-like receptor 1 (ORL-1) is a Gi/o-coupled G-protein-coupled receptor (GPCR) and is the fourth member of the opioid receptor family [10]. ORL-1 is unique in the opioid receptor family since classical endogenous (β-endorphins) and exogenous opioids (morphine) exhibit low or no binding affinity. Instead, ORL-1 is activated by the endogenous neuropeptide nociceptin, which has low affinity for other opioid receptors [47]. ORL-1 is expressed in the primary afferent pain pathway, where it is thought to regulate afferent pain signalling [10, 17, 49], as well as in the hippocampus [25, 48], where it has been shown to regulate spatial learning and memory [44, 55] and memory reconsolidation [53].

It is well documented that various GPCRs can regulate VGCC activity [6, 14, 35, 39, 41, 46, 52, 59] in particular N-type (Cav2.2) calcium channels which are subject to direct membrane delimited inhibition by G protein βγ subunits [26, 30, 31, 37], and L-type calcium channels which are potently augmented by adrenergic receptor mediated activation of protein kinase A (PKA) [19, 34, 46, 51]. ORL-1 has previously been shown to interact with Cav2.2 channels and to produce agonist-independent modulation and co-internalization upon nociceptin treatment [1, 3]. While coupling between Cav2.2 and ORL-1 receptors has been the subject of extensive investigation, much less is known about interactions between Cav1.2 and ORL-1, and investigations into this interaction has yielded mixed results. Knoflach and colleagues demonstrated that acute perfusion with nociceptin inhibited L-type calcium channel currents in hippocampal neurons [43], whereas acute perfusions or chronic treatment in tsA-201 cells did not affect Cav1.2 currents [1, 3].

Considering that Cav1.2 and ORL-1 have overlapping expression in the hippocampus and overlapping function in learning and memory [9, 25, 44, 45, 48, 55], exploring this potential interaction is important. Here, we examined the interplay of Cav1.2 and ORL-1 in tsA-201 cells. We show that ORL-1 independently interacts with both the Cavα1c subunit as well Cavα2δ-1. Moreover, this interaction decreases Cav1.2 peak current density in an agonist-independent manner and reduces plasma membrane expression by disrupting forward trafficking independently of receptor activation.

## Materials and Methods

### Cell Culture and Transfection

All experiments were conducted in tsA-201 cells transfected with cDNA plasmids, mouse Cavα1c (in pcDNA6: GenBank: NM_1159534.2) or rat Cavα1c-BBS (in pMT2: GenBank: M67515.1), rat Cavβ1b (in pcDNA3.1: GenBank: NM_015346.1), Cavα2δ-1 (in pcDNA3.1: GenBank: MP_037051.2), pcDNA3.1HisB, HisORL-1 (in pcDNA3.1: GenBank: NM_000913), and GFP (in pEGFP-N1). For electrophysiology and co-immunoprecipitations, cells were transfected with 3 μg of each cDNA, except GFP, for which 0.5 μg was used (electrophysiology only). For trafficking assays, 2 μg of total cDNA at a 3:2:2:2 ratio (α1c-BBS: α2δ-1: β1b: HisORL-1) was used per 35mm glass-bottomed dish (MatTek Corp., Ashland, MA). Cell transfection using the calcium phosphate method was previously described by Zhang and colleagues [65].

### Electrophysiology

Following transfection, tsA-201 cells were incubated for 24 hours at 37 °C, after which the cell medium was replaced. They were then transferred to a 30 °C incubator for an additional 48 hours. All electrophysiological recordings were conducted at room temperature (22-24°C) using a whole-cell voltage-clamp configuration, an Axopatch 200B, and pClamp 11.2 software. For current-voltage relationship recordings and assessment of calcium-dependent inactivation (CDI), a P/4 subtraction protocol was used to correct for the linear leak component, while the linear leak component was manually corrected for acute perfusions and steady-state inactivation (SSI). For all recordings, traces were digitized at 1kHz then filtered with a Lowpass Gaussian Filter (-3dB cutoff, 1kHz) in Clampfit 11.4.3. Patch pipettes (Sutter Instrument Co., Novato, CA, BF150-86-7.5) were pulled to give a resistance between 2-5 MΩ when filled with a cesium-based internal solution (in mM): 130 CsCl, 2.5 MgCl_2_, 5 EGTA, 10 HEPES, 3 ATP, 0.5 GTP. For IV, SSI, and acute perfusions, coverslips were transferred from the incubated dish to circular Petri dishes (35mm × 10mm) and bathed in external solution (in mM): 10 BaCl_2_, 1 MgCl_2_, 10 HEPES, 10 Glucose, 125 CsCl. For CDI, the following external solution was used (in mM): 110 CsCl, 20 CaCl_2_, 1 MgCl_2_, 10 HEPES, 10 Glucose. The current-voltage (IV) relationship was established using 250ms steps, starting at -90 mV and increasing in 5 mV increments to +50 mV. All data were then fitted with the modified Boltzmann equation:

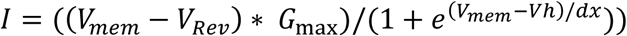

Where V_mem_ represents the membrane potential, G_max_ denotes the maximum conductance, dx is the slope factor, V_h_ is the half-activation voltage, and V_rev_ is the reversal potential. SSI curves were established with a 5s holding potential starting at -110 mV and stepping to +30 mV in 10 mV increments, followed by a test pulse to 0 mV for 140ms. SSI traces were then fit to the following Boltzmann equation:

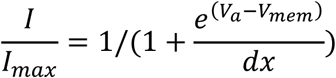

Where V_a_ is the half-inactivation voltage, dx is the slope factor, and V_mem_ represents the membrane potential. CDI was ascertained by running the above IV protocol in the presence of 20 mM CaCl_2,_ followed by fitting the decay of the traces from 0 mV to 30 mV to a single-term exponential function in Clampfit 11.4.3:

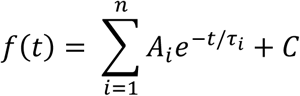

For acute perfusions, tsA-201 cells co-expressing Cav1.2 and ORL-1 were perfused with external solution (VEH) for 1-4 min or until current stabilized, followed by perfusion with 1 μM Nociceptin dissolved in external solution for 4-7 mins.

### Co-immunoprecipitations

Twenty-four hours after transfection, the tsA-201 cells were treated with fresh medium and incubated at 37 °C for an additional 48 hours. After washing with PBS : (in mM) 137 NaCl, 2.7 KCl, 10 Na_2_HPO_4_-7H_2_O, 2 KH_2_PO_4_, cells were lysed with channel lysis buffer (in mM): 50 Tris-HCl, pH 7.5, 150 NaCl, 1% Igepal, 1% Triton X-100, 0.2% SDS, 5 EDTA + protease inhibitor (Complete, Mini, EDTA-free Protease Inhibitor Cocktail, Roche). Lysates were extracted and then centrifuged for 15 minutes at 13,300 rpm at 4 °C. Protein content was then quantified using 1mg/mL BSA standards and a Jenway Spectrophotometer. The volume equivalent to 1-2mg of protein was then incubated with anti-Xpress monoclonal antibody (Invitrogen) or anti-Cav1.2 (CACNA1C) antibody (Alomone CAT# ACC-003) at a ratio of 1-2 μg antibody per 1mg of protein. The following day, the lysates were incubated with 50 μl of Protein A or Protein G Sepharose 4 Fast Flow (Millipore Sigma) for 1.5 hours. Following three washes with lysis buffer, 32 μL of Laemmli buffer (6% SDS, 187.5mM, 30% Glycerol, 0.0015% Bromophenol Blue, 7.5% 2-mercaptoethanol) was used to denature proteins.

### Western Blots

For experiments involving immunoprecipitation of ORL-1 followed by immunoblotting for Cavα1c, samples were heated in a 55 °C hot bath for 10 minutes before being loaded into a lab-made 10% SDS-PAGE gel. For Cavα1c immunoprecipitation and ORL-1 immunoblotting, samples were left at room temperature for 30 minutes before being loaded into a lab-made 7.5% SDS-PAGE gel. Membranes were then blocked for 1 hour with 5% skim milk before being incubated overnight with 1:500 Nociceptin Receptor (OPRL1) Polyclonal Antibody or 1:250 Anti-Cav1.2 (CACNA1C) Antibody (Alomone Labs). The following day, cells were washed with TBST (in mM): 2 Tris, 137 NaCl, 10ml Tween, then incubated for 1 hour with 1:5000 anti-rabbit HRP secondary antibody (Jackson Immunoresearch) or 1:5000 anti-mouse HRP secondary antibody (Jackson Immunoresearch). Membranes were then scanned using a C-DiGit blot scanner (LI-COR) and Imagine Studio Digits Ver 5.2.

### Trafficking Assays and Confocal Imaging

Prior to transfection, glass-bottomed dishes (MatTek Corp., Ashland, MA) were precoated with poly-ʟ-lysine. After transfection and 3-day incubation at 37 °C, cells were washed 2-3 times with Krebs–Ringer solution with HEPES (KRH) (in mM): 125 NaCl, 5 KCl, 1.1 MgCl2, 1.2 KH2PO4, 2 CaCl2, 6 Glucose, 25 HEPES,1 NaHCO3, adjusted to pH 7.4. For experiments assessing endocytosis kinetics, cells were incubated with 10 μg/ml α-bungarotoxin Alexa Fluor® 594 conjugate (BTX594) (Thermo Fisher Scientific) at 17°C for 30 minutes. Cells were washed with KRH to remove excess BTX594, then incubated in KRH at 37°C for 0, 5, 10, 20, and 30 minutes. Cells were washed with PBS before being fixed with 4% PFA and 4% Sucrose in PBS for 5 minutes. Cells were then permeabilized with 0.05% Triton X-100 in PBS for 10 minutes, followed by blocking with 5% BSA and 20% GS in PBS for at least 45 minutes. Following blocking, cells were incubated for 1 hour with 1:200 anti-CaV1.2 (Alomone Labs) in PBS containing 5% BSA and 20% Goat Serum. Cells were then washed and incubated for 1 hour with 1:500 secondary conjugated Ab Anti-rabbit AF488 (Donkey anti-Rabbit Alexa Fluor^TM^ Plus, Thermo Fisher Scientific). Blocking and antibody incubation were performed at room temperature (22-24°C). For experiments examining forward trafficking, cells were incubated with 10 μg/ml unlabelled BTX for 30 minutes at 17°C to saturate the plasma membrane Cav1.2-BBS binding sites. Cells were washed with KRH and then incubated with 10 μg/ml BTX594 for 0, 5, 10, 20, and 30 minutes. Cells were then fixed and permeabilized as described for endocytosis experiments. Cells were covered with SlowFadeTM Gold Antifade Mountant (Thermo Fisher Scientific) and stored in the dark overnight at 3 °C. On the following day, cells were examined with a Leica SP8 confocal microscope using a 63×/1.4 numerical aperture oil-immersion objective in 16-bit mode. For endocytosis experiments, cells at time point T = 0 minutes (T0) were used to calibrate acquisition settings for AF594 and AF488 to avoid oversaturation, while cells at time point T = 30 minutes (T30) were used to calibrate forward trafficking experiments.

Acquisition settings were then kept constant for the rest of the experiment. Confocal images were processed in ImageJ, and the region-of-interest data were then transferred to GraphPad Prism. Endocytosis kinetics were determined by fitting data to the following one-phase decay equation:

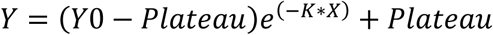

Where X is time, Y0 is the Y value at time zero, K is the rate constant, and plateau is the Y value when time equals infinity.

### Statistical analysis

All data are presented as means plus standard error of the mean. All data were tested for outliers in GraphPad Prism, followed by a normality test. Data sets deemed normal were analyzed with an unpaired Welch’s t-test. For data that failed the normality test, differences were analyzed using an unpaired Mann-Whitney t-test. For acute perfusions, GraphPad Prism’s paired, normal (Gaussian) distribution t-test was used. For all statistical analyses, significance was set at 0.05. Asterisks are used to denote significance: *p <0.05, ****p < 0.0001.

## Results

### ORL-1 and Cav1.2 interact in tsA-201 cells

We previously showed that ORL-1 co-immunoprecipitates with Cav2.2 in tsA-201 cells and in dorsal root ganglion neurons [3]. Thus, to ascertain whether ORL-1 forms a similar complex with Cav1.2, co-immunoprecipitations were conducted in tsA-201 cells co-expressing Cav1.2 (Cavα1c, Cavβ1b and Cavα2δ-1) and His-tagged ORL-1. Immunoprecipitation with an anti-Xpress antibody followed by immunoblotting with an anti-Cavα1c antibody revealed that Cavα1c co-immunoprecipitated with ORL-1 (**Figure 1a**). To verify this interaction, we immunoprecipitated Cavα1c followed by immunoblotting with an anti-ORL-1 antibody. As shown in **Figure 1b**, ORL-1 was detected following immunoprecipitation of Cavα1c. Co-immunoprecipitations were observed when the pore-forming subunit was co-expressed with Cavβ1b and Cavα2δ-1, or when expressed alone, indicating that the Cavβ1b and Cavα2δ-1 subunits were not required for the formation of receptor-channel complexes. Given that the Cavα2δ-1 subunit interacts with several other membrane proteins besides calcium channels [15, 65], a separate set of experiments was conducted to explore whether Cavα2δ-1 interacts with ORL-1 independently of Cavα1c. As seen in **Figure 1c**, Cavα2δ-1 co-immunoprecipitated with ORL-1 when expressed in the absence of other calcium channel subunits. Together, these results demonstrate that ORL-1 interacts separately with two components of the Cav1.2 channel complexes.

**Figure 1.**
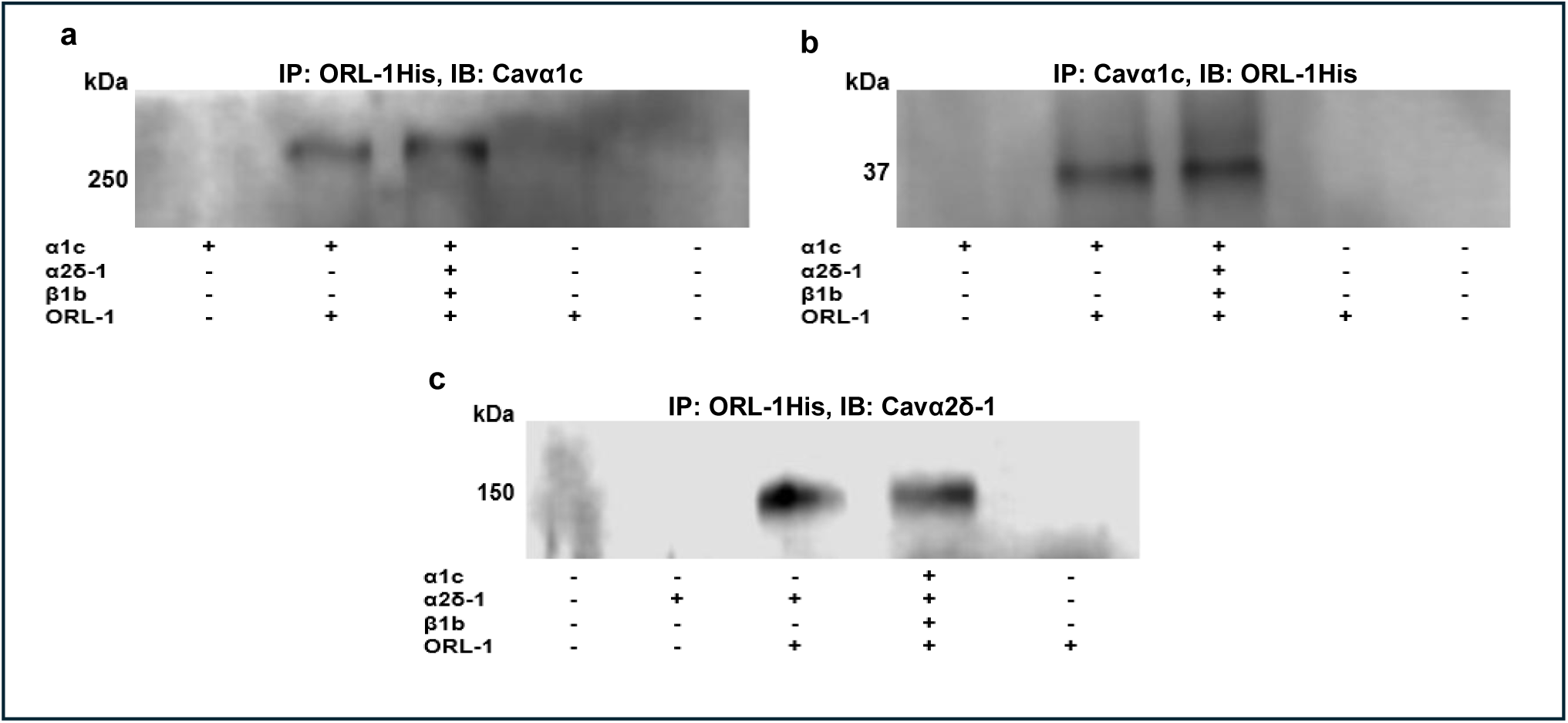
ORL-1 interacts with Cavα1c and Cavα2δ-1 of the Cav1.2 channel complex. **a**) Co-immunoprecipitation from tsA-201 cell lysate, with immunoprecipitation for ORL-1-HIs and immunoblotting for Cavα1c (n = 5). Cavα1c co-immunoprecipitates with ORL-1 regardless of auxiliary subunit expression. **b**) Immunoprecipitation for Cavα1c, with immunoblotting for ORL-1, demonstrating ORL-1 co-immunoprecipitates with Cavα1c in the presence and absence of auxiliary subunits (n = 3). **c**) Immunoprecipitation for ORL-1-His, with immunoblotting for Cavα2δ-1, demonstrating that Cavα2δ-1 co-immunoprecipitates with ORL-1 when co-expressed both in the presence and absence of Cavα1c (n = 3).

### ORL-1 decreases Cav1.2 current density without modulating its biophysical properties

Our prior work revealed that interactions between ORL-1 and Cav2.2 have functional consequences on Cav2.2 activity, even in the absence of ORL-1 activation [1, 3]. Thus, given the formation of a Cav1.2-ORL-1 protein complex, we wanted to test whether this interaction may alter Cav1.2 activity. To address this, Cav1.2 was expressed in tsA-201 cells with or without ORL-1. Whole-cell patch- clamp recordings were then used to assess current-voltage relationship, steady-state inactivation, and CDI. **Figure 2a** depicts families of whole-cell current traces for Cav1.2 in the presence and absence of ORL-1; **Figure 2b** shows the current-voltage relationships of Cav1.2 with and without ORL-1 in 10 mM Ba^2+^. The IV curve suggests a small shift in half-activation voltage. However, this trend did not reach statistical significance (**Figure 2c**). The current-voltage relations shown in **Figure 2c** include only cells with sufficiently large currents that could be fitted appropriately with the Boltzmann equation, leading to an underestimation of the effect of ORL-1 on the overall Cav1.2 peak current density. When cells with small currents were included in the peak current density analysis, it emerged that ORL-1 significantly decreased Cav1.2 peak currents (**Figure 2d**). Peak current density was determined by selecting the peak current from each individual trace. ORL-1 did not affect Cav1.2 steady-inactivation properties (**Figure 3a**) or CDI tested in 20 mM Ca^2+^ to trigger robust calcium elevations (**Figure 3b**). As a negative control, Cav3.2 was co-expressed with ORL-1 in tsA-201 cells. As expected, based on our group’s previous findings, ORL-1 did not alter Cav3.2 activity (**Supplementary Figure 1 and Figure 2a**). Together, these results demonstrate that the interaction between ORL-1 and Cav1.2 has no significant effects on the biophysical properties of Cav1.2 channels but reduces Cav1.2 current amplitude. This effect could, in principle, be due to reduced single-channel conductance, reduced maximum open probability, or an effect on the number of channels in the plasma membrane.

**Figure 2.**
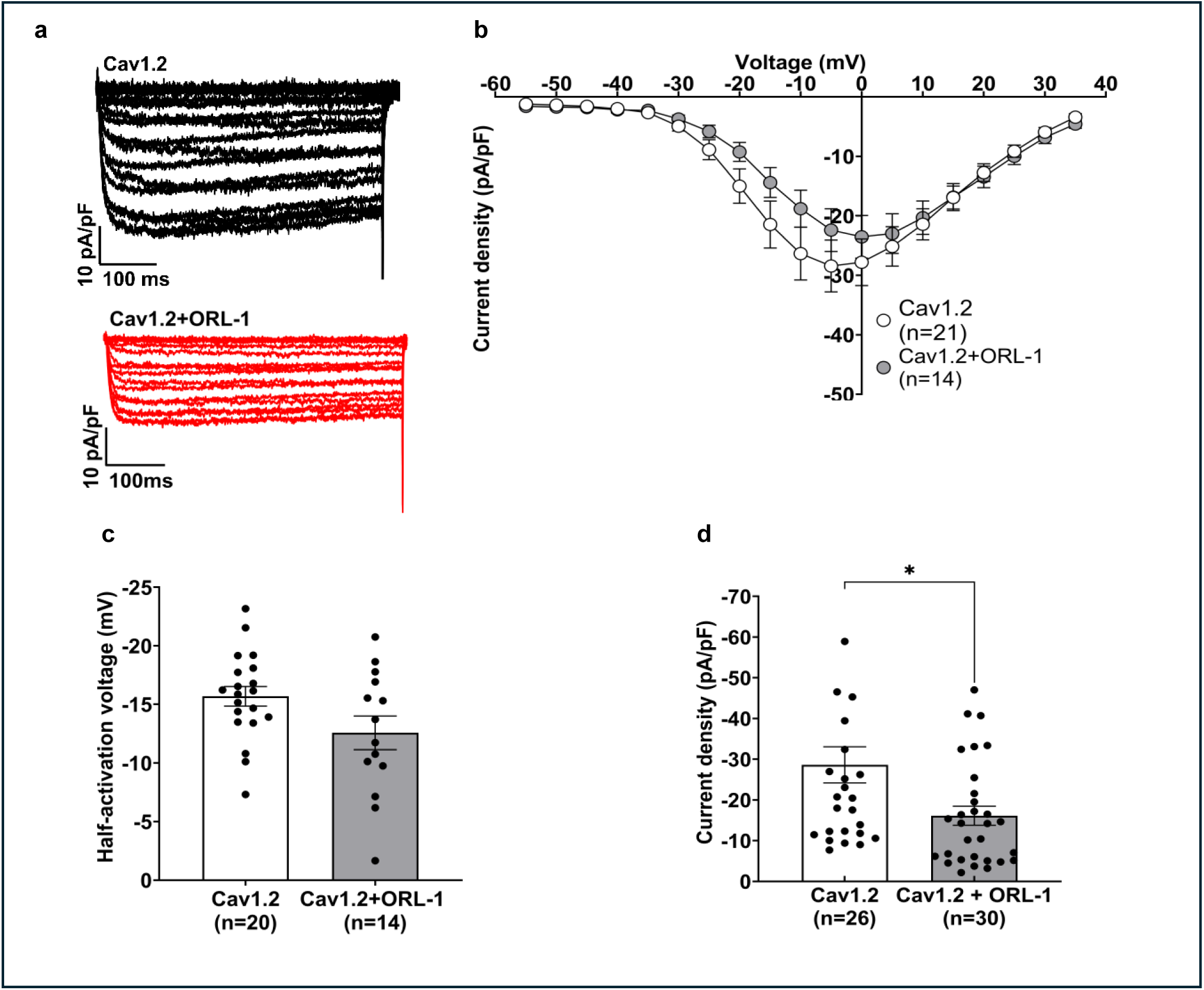
Co-expression of ORL-1 and Cav1.2 in tsA-201 cells decreases Cav1.2 peak current density without affecting biophysical properties. **a)** Representative whole cell traces for Cav1.2 and Cav1.2+ORL-1. Traces were evoked by depolarizing steps from -90 mV to +50 mV in 5 mV increments. **b)** Current-voltage relationships for Cav1.2 co-expressed with or without ORL-1 (n=21, n=14). All currents were normalized by cell capacitance. **c)** Half-activation voltage of Cav1.2 in the presence and the absence of ORL-1. **d)** Peak current density for Cav1.2 co-expressed with or without ORL-1. ORL-1 significantly reduced Cav1.2 peak current density (p = 0.0170, n=26, n=30). Peak current densities were taken from each individual trace, a majority of which occurred between -10 mV and 10 mV. Data are expressed as mean ± SEM.

**Figure 3.**
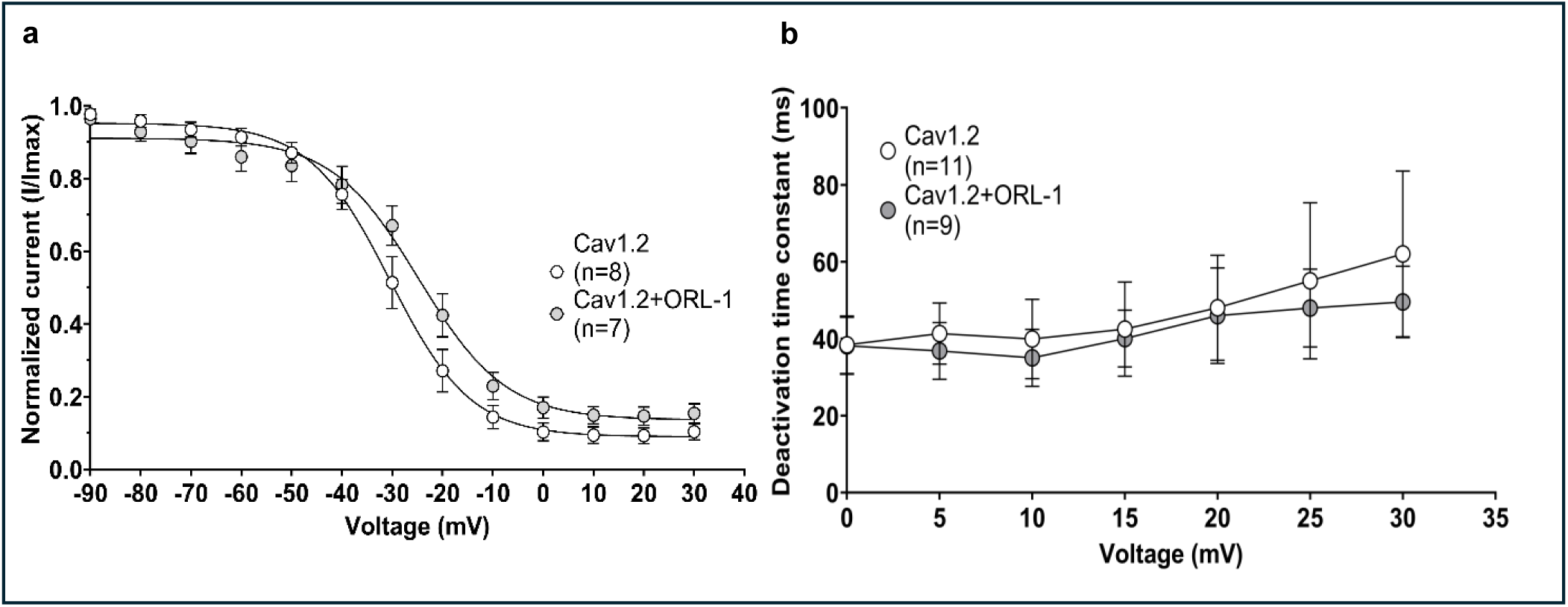
Co-expression of ORL-1 and Cav1.2 does not alter Cav1.2 voltage dependence of inactivation or CDI. **a)** Steady-state inactivation curves for Cav1.2 co-expressed with or without ORL-1 in tsA-201 cells. ORL-1 did not significantly alter the slope or half-inactivation voltage. **b)** _2+_ Calcium-dependent inactivation of Cav1.2 expressed with or without ORL-1 in 20 mM Ca^2+^ external solutions, shown in the form of time constants of inactivation. Voltages around the average peak-current voltage (15 mV in 20 mM Ca^2+^) were fitted with a single exponential. At all tested voltages, there was no difference between Cav1.2 alone or co-expressed with ORL-1. Data are expressed as mean ± SEM.

### Activation of ORL-1 has no effect on Cav1.2 currents

Activation of ORL-1 has been shown to mediate potent inhibition of Cav2.2 channels in neurons and in tsA-201 cells via Gβγ [1, 3]. In contrast, GPCR modulation of L-type channels often involves activation or inhibition of protein kinase A, which acts either directly on the channel [34, 51] or indirectly modulates current activity via phosphorylation of the small GTPase Rad [50]. To ascertain whether activation of ORL-1 alters Cav1.2 channel activity, tsA-201 co-expressing Cav1.2 and ORL-1 were perfused with external solution as a vehicle (VEH) for 1-4 mins, followed by 1 µM of nociceptin for 4-7 mins. Throughout the perfusion, Cav1.2 currents were evoked by depolarizing steps to 0mV every 20s. As shown in Figure 4, treatment with nociceptin did not affect Cav1.2 activity compared with vehicle. Similarly, ORL-1 activation with nociceptin did not affect Cav3.2 activity (**Supplementary Figure 2b**). These results are consistent with acute perfusion results reported in a previous report from our lab and add to the growing body of literature suggesting that ORL-1 agonist-mediated activation has no effect on Cav1.2 [3].

**Figure 4.**
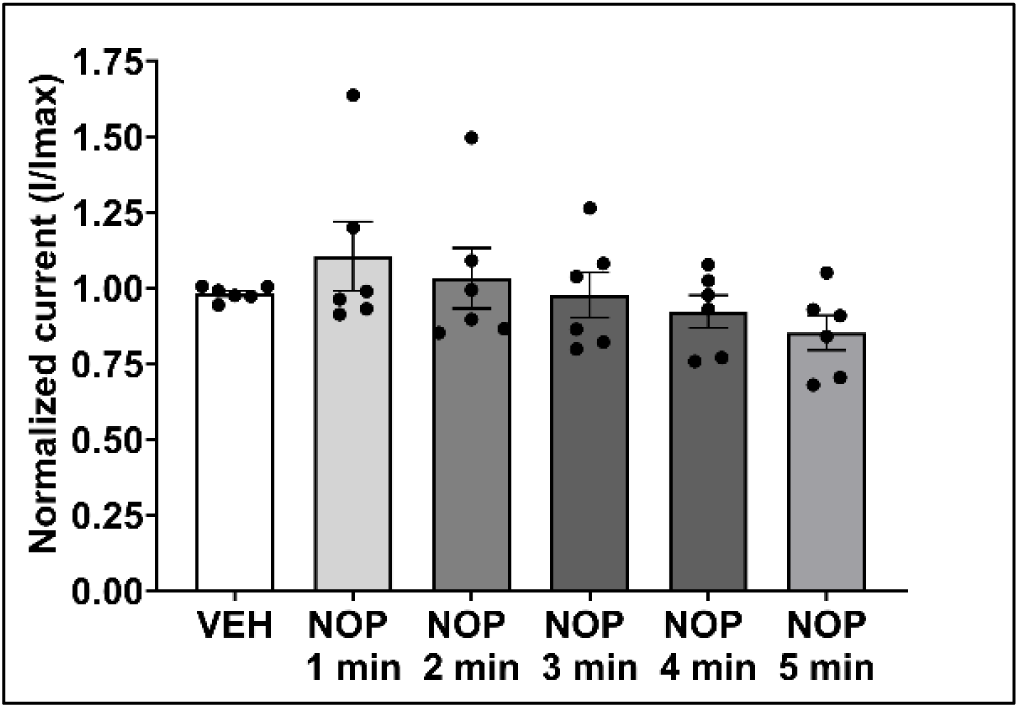
Effect of nociceptin on Cav1.2 current amplitude. ORL-1 and Cav1.2 were co-expressed in tsA-201 cells and treated with acute continuous perfusion of external solution (VEH), followed by 1 µM nociceptin dissolved in external solution. Cav1.2 currents were elicited with voltage steps to 0mV every 20s. Activation of ORL-1 with nociceptin had no effect on Cav1.2 currents (n = 6). Data are expressed as mean ± SEM.

### ORL-1 decreases Cav1.2 membrane expression by disrupting forward trafficking

The ability of ORL-1 to decrease Cav1.2 current density through simple co-expression suggests that it either produces a tonic agonist-independent inhibition or alternatively perhaps disrupts membrane trafficking. To explore Cav1.2 membrane trafficking, we used a Cavα1c subunit with α-bungarotoxin binding sites inserted in the extracellular loop between S5 and the P loop of domain II, as previously designed by us [24]. TsA-201 cells expressing the Cav1.2-BBS construct were incubated with an α-bungarotoxin Alexa Fluor 594 conjugate (α-BTX594) for 30 mins at 17 °C, followed by incubation at 37 °C for 0, 5, 10, 20 or 30 mins. Cells were then fixed, permeabilized, and stained with an anti-α1c primary and an anti-rabbit AF488 secondary antibody. This allowed for quantification of Cav1.2 membrane staining, normalized to total Cav1.2 expression. **Figure 5a** illustrates membrane BBS staining and total staining at T0, T10 and T30. Endocytosis kinetics were assessed by comparing relative membrane expression normalized to T0. Over five experiments, we found no difference in Cav1.2 endocytosis between co-expression with ORL-1 and expression alone (**Figure 5b**). However, at T0, we found that ORL-1 significantly decreased overall membrane expression of Cav1.2 compared to Cav1.2 alone (**Figure 5c**). This suggests that the effect of ORL-1 on Cav1.2 membrane expression may perhaps be caused by decreased forward trafficking.

**Figure 5.**
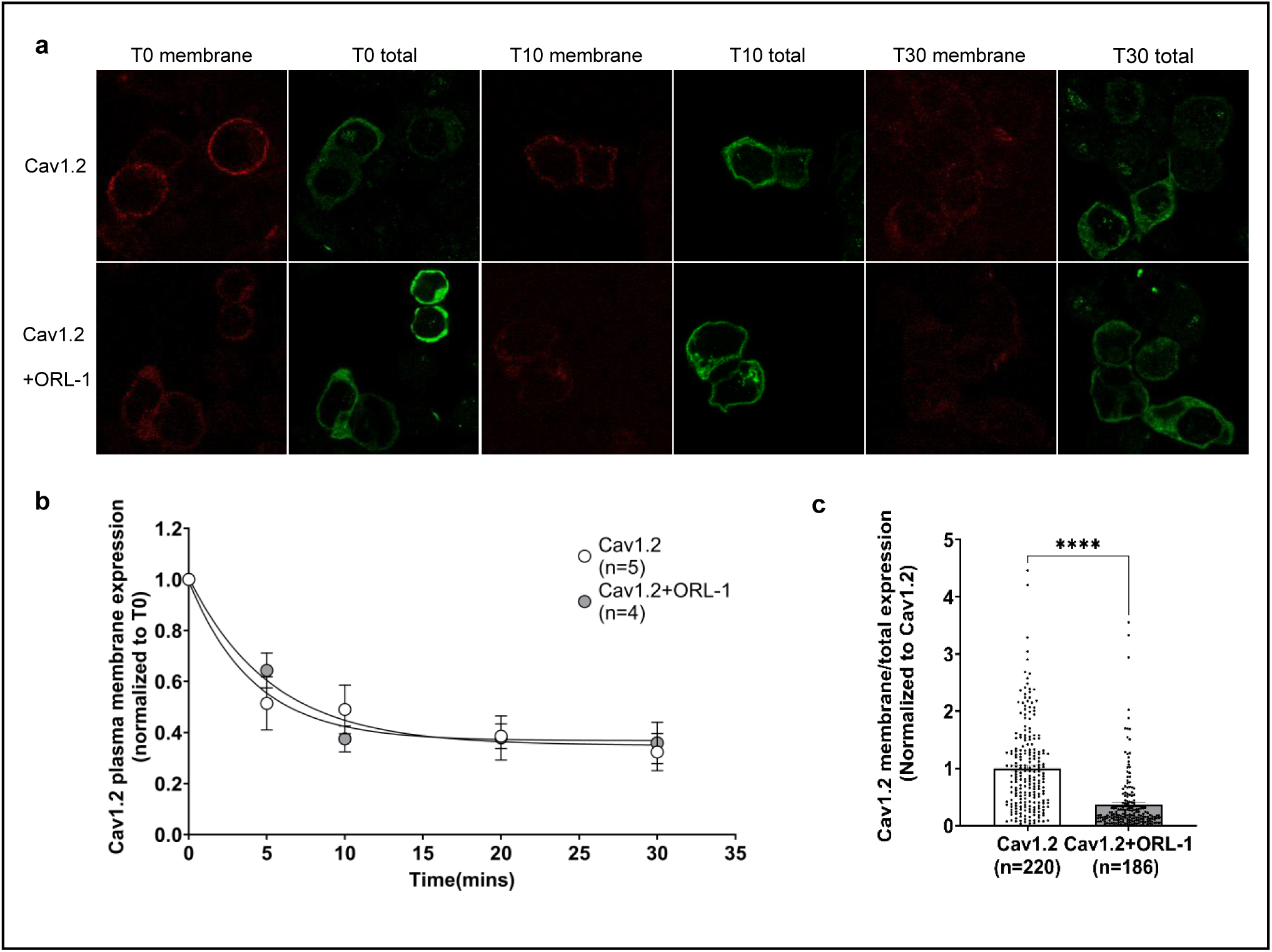
Effect of ORL1 on Cav1.2 trafficking. Cav1.2 membrane expression and endocytosis kinetics were explored by expressing a Cav1.2 containing an α-bungarotoxin binding site with or without ORL-1. Incubation with an α-bungarotoxin AF-594 conjugate at 17 °C for 30 minutes allowed for membrane-specific staining. Cells were then incubated in KRH at 37 °C for 0, 5, 10, 20 and 30 minutes before being fixed, permeabilized, and stained with an anti-α1c primary and anti-rabbit-488 conjugate. **a)** Visual representation of Cav1.2-BBS endocytosis experiments. At T0 (left), there is strong Cav1.2 membrane expression (red) and strong total Cav1.2 expression (green). At T10 (middle), there is a visible decrease in Cav1.2 membrane expression. At T30 (right), Cav1.2 membrane staining has drastically decreased. **b)** Cav1.2 membrane expression at T0, T5, T10, T20 and T30 was normalized to T0 for both Cav1.2 and Cav1.2+ORL-1 and then graphed as relative expression over time and fitted with a one-phase decay equation. ORL-1 did not alter Cav1.2 endocytosis kinetics (n = 5). **c)** Cav1.2 membrane expression is normalized to Cav1.2 total expression, then normalized to the Cav1.2 control condition as 1. The bar graph produced shows that co-expressing Cav1.2 with ORL-1 significantly decreases channel membrane expression compared to Cav1.2 alone (p < 0.0001, n = 220, n = 186). Data are expressed as mean ± SEM.

To investigate forward trafficking, tsA-201 cells were first incubated with unlabelled α-bungarotoxin to saturate all Cav1.2-BBS binding sites. TsA-201 cells were then incubated with α-BTX594 at 37 °C for 0, 5, 10, 20 or 30 minutes. Thus, any newly inserted Cav1.2-BBS channels added to the plasma membrane during incubation at 37 °C become fluorescently labelled (**Figure 6a**). When Cav1.2 membrane expression was normalized to total Cav1.2-BBS expression at each time point, we found that ORL-1 coexpression reduced the insertion of Cav1.2 channels (**Figure 6b**). This is also evident when the data at time point T30 were normalized to the Cav1.2 control condition, revealing that ORL-1 significantly decreased Cav1.2 membrane expression (**Figure 6c**).

**Figure 6.**
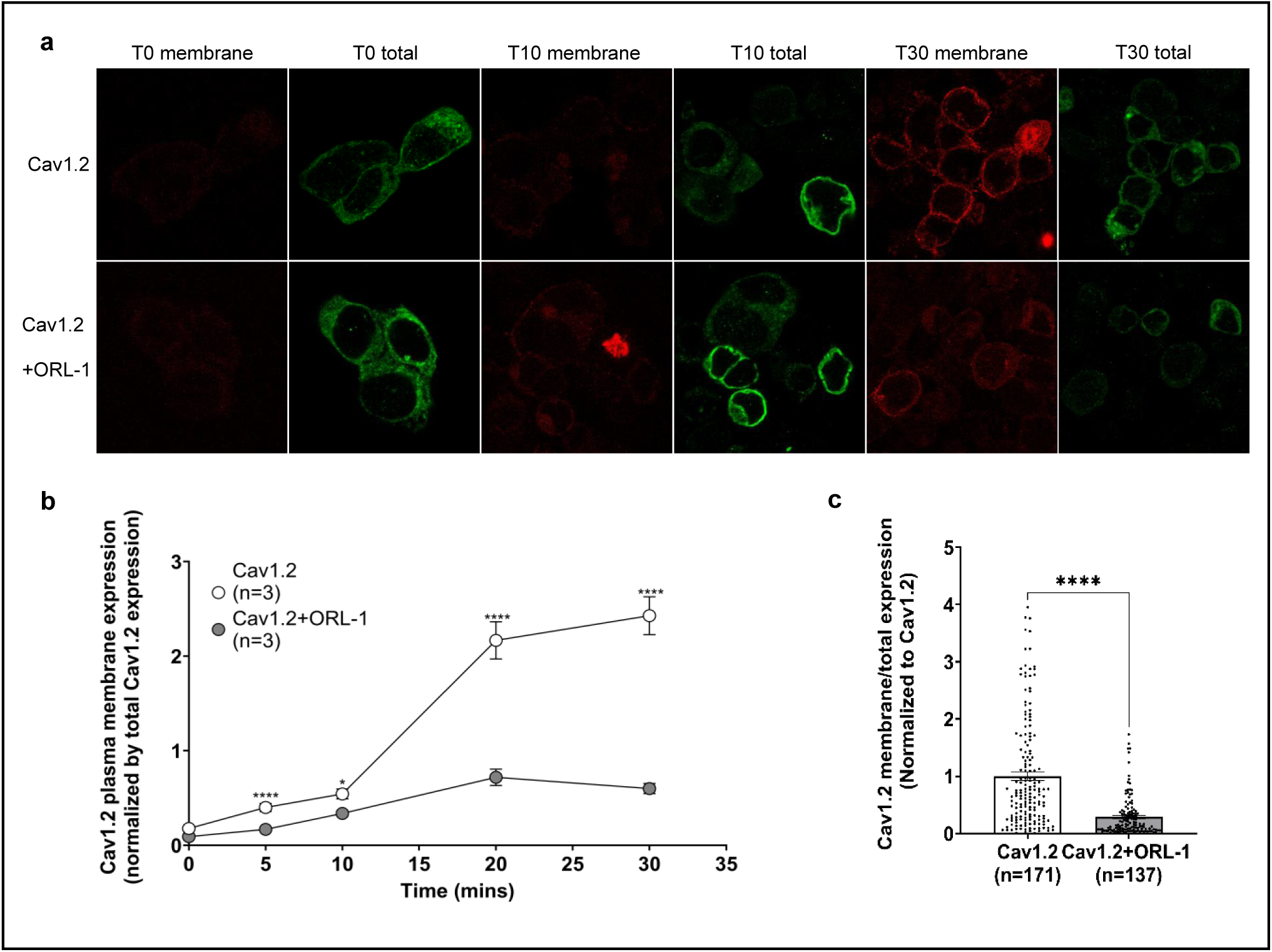
ORL-1 disrupts Cav1.2 forward trafficking. TsA-201 cells expressing Cav1.2-BBS with or without ORL-1 were incubated with an unlabelled α-bungarotoxin at 17 °C for 30 minutes to saturate membrane BBS binding sites before being incubated at 37 °C with α-bungarotoxin AF-594 for 0, 5, 10, 20 and 30 minutes. Only newly inserted membrane Cav1.2 channels during this time were therefore labelled with AF594. **a)** Visual representation of the forward trafficking experiments. At T0 (left), Cav1.2 membrane expression (red) is virtually nonexistent while total Cav1.2 expression (green) is strong. At T10 (middle), membrane staining has increased relative to T0 but is still minimal. At T30 (right), membrane expression has increased drastically compared to T0 and T10. **b)** Cav1.2 membrane expression in each cell, normalized to its own total expression for every time point. ORL-1 decreases Cav1.2 membrane expression starting at T5, and with the largest difference observed at T20 and T30. T0 (p < 0.0001, n = 87, n = 104), T10 (p = 0.0364, n = 122, n = 112, T20 p < 0.0001, n = 159, n = 113, T30 p < 0.0001, n = 171, n = 137. **c)** Cav1.2 membrane expression was normalized by total expression, then normalized to Cav1.2 control for each experiment. ORL-1 significantly decreases Cav1.2 membrane expression (p < 0.0001, n = 171, n = 137). Data are expressed as mean ± SEM.

Together, the endocytosis and forward trafficking experiments strongly suggest that ORL-1 decreases Cav1.2 membrane expression by disrupting forward trafficking, consistent with the overall reduction in whole-cell current densities observed in our electrophysiological studies.

## Discussion

GPCR modulation of VGCC activity is well-documented and occurs in various forms [6, 14, 35, 39, 41, 46, 52, 59]. Members of the Cav2 channel family (P/Q, N, R-type) typically undergo membrane-delimited direct modulation by the Gβγ of GPCRs, which can bind to two distinct regions of the I-II linker [7, 18, 61, 64]. In contrast, GPCRs typically modulate L-type calcium channels through protein kinase and phosphatase pathways. A classic example is the Cav1.2- β-adrenergic receptor (βAR) signalling complex. In neurons, Cav1.2, βAR2, and PKA are tightly associated, enabling rapid potentiation of Cav1.2 currents via PKA-mediated phosphorylation [19, 46, 51]. Similarly, Gi/o-coupled CB1 and GABA B receptors can inhibit LTCC by inhibiting the AC**→**cAMP**→**PKA pathway [2, 32].

In the present study, we report a distinct mechanism by which GPCRs modulate Cav1.2 channel activity, in which ORL-1 independently interacts with the Cavα1c and Cavα2δ-1 subunits of the Cav1.2 channel complex. This interaction decreases Cav1.2 membrane expression and peak current density through disrupting forward trafficking. The observed decrease in Cav1.2 peak current density could, in principle, be explained by constitutive ORL-1 activity [3] that may suppress PKA activation, thereby decreasing Cav1.2 phosphorylation. However, unless such constitutive activity were to occur at saturating levels, further treatment with nociceptin should have amplified such inhibition, which was not observed. Furthermore, if inhibition of PKA-mediated phosphorylation were exclusively responsible for the decrease in peak current density, then one would have expected to observe similar Cav1.2 membrane expression in the presence and the absence of ORL-1. Instead, our confocal microscopy experiments demonstrate a substantial decrease in Cav1.2 membrane expression and a disruption in forward trafficking when co-expressed with ORL-1.

The independent interaction between ORL-1 and Cavα2δ-1 may potentially account for the observed decrease in membrane expression and the disruption of forward trafficking. We have previously reported that large-conductance Ca^2+^ and voltage-dependent K^+^ channels (BK) can interact with Cavα2δ-1 and decrease high-voltage activated channel membrane expression and peak current density, including that of Cav1.2 [65]. Considering that Cavα2δ-1 plays a crucial role in membrane trafficking [11, 33, 63], the effect observed in [65] is likely mediated by BK channels competing Cavα2δ-1 subunits away from the channels, thereby disrupting their membrane trafficking. ORL-1 receptors may similarly compete away Cavα2δ-1, thereby decreasing Cav1.2 membrane expression via reduced forward trafficking. In our previous work on Cav2.2 interactions with ORL-1 receptors, we did not observe a suppression of forward trafficking [1] even though Cavα2δ-1 is also required for the membrane trafficking of Cav2.2, which would argue against the competition hypothesis. However, it is possible that Cav1.2 and Cav2.2 channels interact with separate pools of Cavα2δ-1 subunits such that ORL-1 might selectively sequester a Cav1.2-associated pool, thus reducing membrane expression. Alternatively, a direct interaction between ORL-1 and the Cav1.2α1c subunit may directly hinder forward trafficking of the channel. At this point, we cannot distinguish among these alternatives.

In the central and peripheral nervous systems, the interaction between ORL-1 and Cav1.2 may have important implications for neuronal function. It has been reported that ORL-1 expression is increased in neuropathic pain [8, 16, 37] while selective knockdown of Cav1.2 in the spinal cord temporarily attenuates mechanical allodynia and neuronal hyperexcitability in spinal-nerve ligated rats [23, 27]. An ORL-1 receptor-mediated inhibition of Cav1.2 membrane expression may thus serve a protective analgesic function. Similarly, in the hippocampus, it is well established that ORL-1 and Cav1.2 have overlapping functional implications in learning and memory [9, 25, 48, 56]. It is thus possible that alterations in ORL-1 expression could affect synaptic plasticity by regulating Cav1.2 channel density in the plasma membrane. Further studies in native systems will be required to test such concepts, and we acknowledge that expression studies in tsA-201 cells have inherent limitations. That being said, our findings have uncovered a novel complex between a G protein-coupled receptor and Cav1.2 channels. Moreover, we note that our data add to a growing body of literature showing that Cavα2δ subunits are highly promiscuous – acting not merely as calcium channel subunits, but also as interactors of BK potassium channel [65], NMDA receptors [15, 66] and AMPA receptors [36], and now ORL-1, all of which are expressed in the primary afferent pain pathway. Given that Cavα2δ subunits are the key analgesic target for gabapentinoids, these multi-target interactions may have important implications for understanding how gabapentinoids suppress pain signalling. Future studies examining whether Cavα2δ regulates ORL-1 receptor function and activity, and whether gabapentinoids alter this modulation, will be insightful.

## Abbreviations

VGCCs: Voltage-gated calcium channels
GPCRs: G-protein-coupled receptors
ORL-1: Opioid-like receptor 1

## Acknowledgements

We thank Lina Chen for providing technical support.

## Code availability

Not Applicable.

## Author contributions

Data acquisition: A.J.S., I.A.S. Methodology: A.J.S., I.A.S., L.F., M.A.G., G.W.Z. Writing-original draft preparation: A.J.S., writing – review and editing: A.J.S., I.A.S., L.F., M.A.G., G.W.Z. Project administration: G.W.Z. Funding Acquisition: G.W.Z. All authors have read and agreed to the published version of the manuscript.

## Data availability

Data published in this study are available from the authors upon reasonable request

## Ethics approval

Not applicable

## Consent to participate

Not applicable

## Consent for publications

Not applicable

