## Supplementary Information for "Effect of ORL-1 on Cav1.2 calcium channels"

For the article

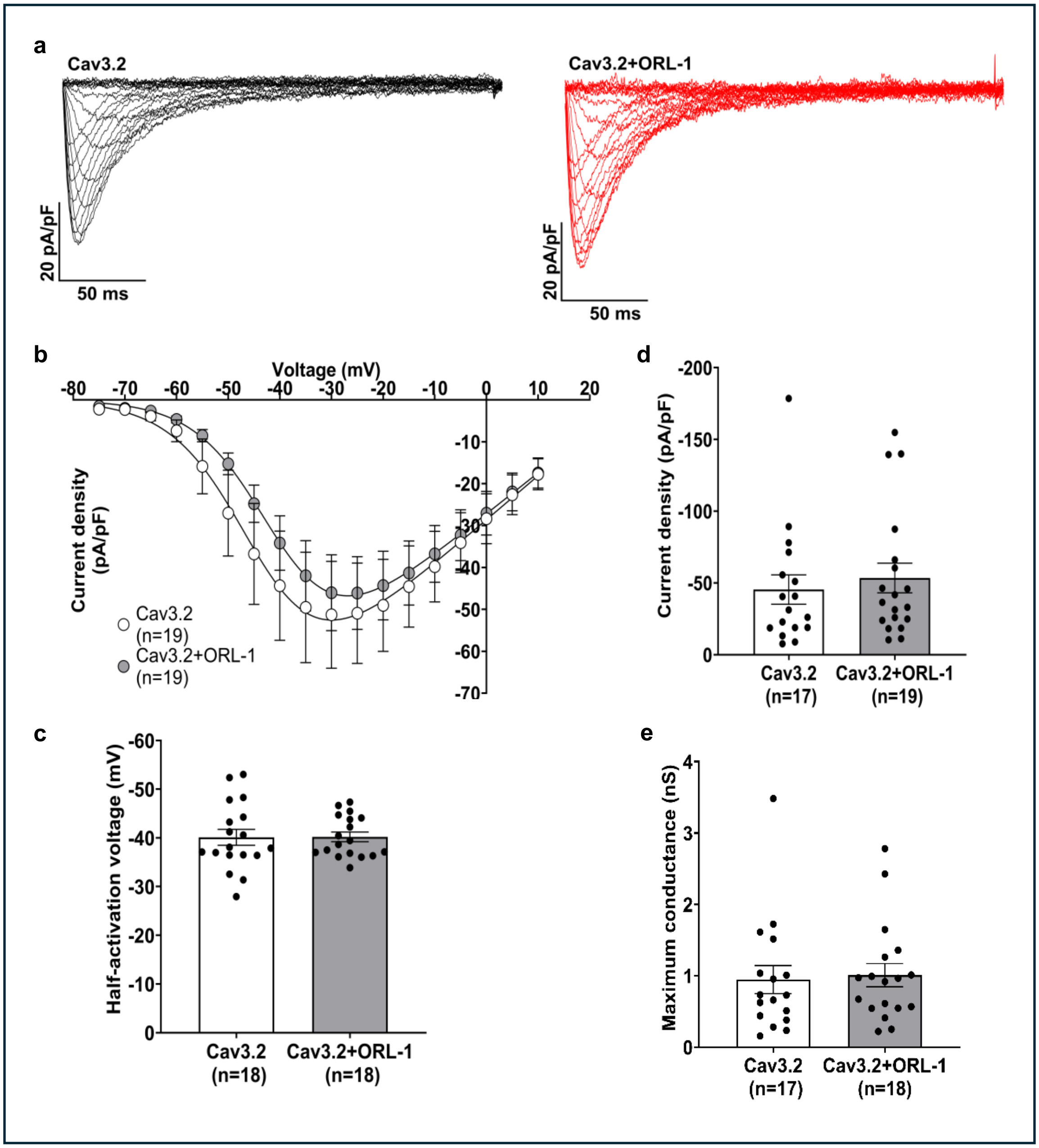


**Supplementary Figure 1** Co-expression of ORL-1 and Cav3.2 in tsA-201 cells does not alter Cav3.2 peak current density or other biophysical properties. **a)** Representative whole cell traces for Cav3.2 and Cav3.2+ORL-1. Currents were evoked by depolarizing steps from -90 mV to +10 mV in increments of 5 mV. **b)** Current-voltage relationship for Cav3.2 with or without co-expression of ORL-1 (n = 19, n = 19). All currents were normalized by cell capacitance. **c)** Peak current density for Cav3.2 co-expressed with or without ORL-1. The presence of ORL-1 did not alter Cav3.2 peak current. **d)** Half activation voltage for Cav3.2 in the presence and absence of ORL-1. **b)** Co-expression of ORL-1 and Cav3.2 did not alter Cav3.2 maximum conductance (Gmax). Data are expressed as mean ± SEM.

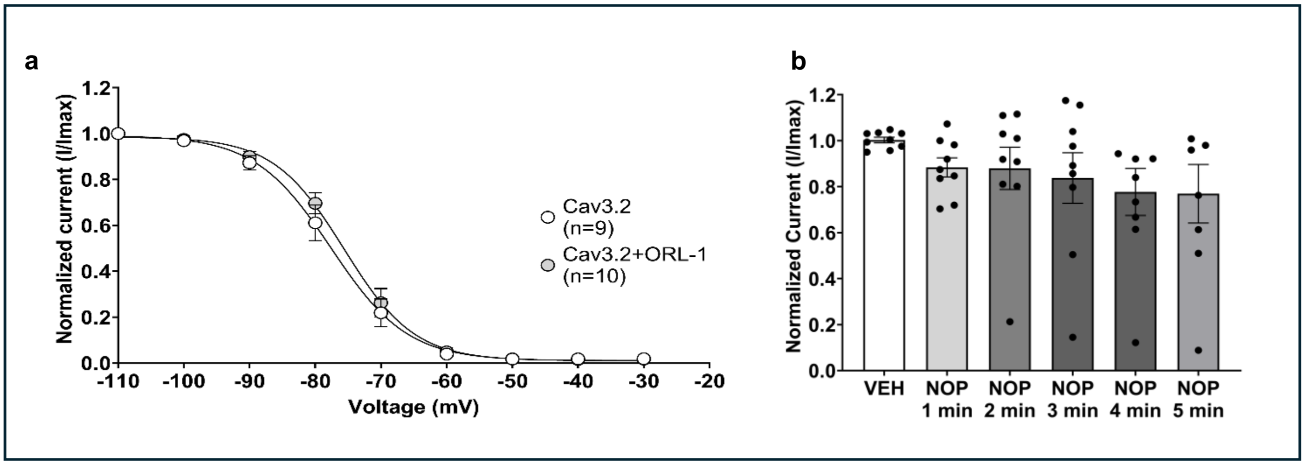


**Supplementary Figure 2 a)** Steady state inactivation curves for Cav3.2 with and without co-expression of ORL-1. Co-expression of ORL-1 with Cav3.2 does not affect voltage-dependent inactivation. **b)** Effect of nociceptin on Cav3.2 current amplitude. ORL-1 and Cav3.2 were co-expressed in tsA-201 cells and treated with acute continuous perfusion of external solution (VEH), followed by 1 μM nociceptin dissolved in external solution. Cav3.2 currents were elicited with voltage steps to -30 mV every 20s. Activation of ORL-1 with nociceptin had no effect on Cav3.2 currents (n = 6). Data are expressed as mean ± SEM.
